# Explaining face representation in the primate brain using different computational models

**DOI:** 10.1101/2020.06.07.111930

**Authors:** Le Chang, Bernhard Egger, Thomas Vetter, Doris Y. Tsao

## Abstract

Understanding how the brain represents the identity of complex objects is a central challenge of visual neuroscience. The principles governing object processing have been extensively studied in the macaque face patch system, a sub-network of inferotemporal (IT) cortex specialized for face processing. A previous study reported that single face patch neurons encode axes of a generative model called the “active appearance” model, which transforms 50-d feature vectors separately representing facial shape and facial texture into facial images. However, a systematic investigation comparing this model to other computational models, especially convolutional neural network models that have shown success in explaining neural responses in the ventral visual stream, has been lacking. Here, we recorded responses of cells in the most anterior face patch AM to a large set of real face images and compared a large number of models for explaining neural responses. We found that the active appearance model better explained responses than any other model except CORnet-Z, a feedforward deep neural network trained on general object classification to classify non-face images, whose performance it tied on some face image sets and exceeded on others. Surprisingly, deep neural networks trained specifically on facial identification did not explain neural responses well. A major reason is that units in the network, unlike neurons, are less modulated by face-related factors unrelated to facial identification such as illumination.

## Introduction

Primates are able to recognize objects invariant to changes in orientation and position. Neurons in macaque face patch AM represent facial identity independent of head orientation (Freiwald and Tsao, 2010), therefore providing a unique opportunity to study how invariant object identity is represented in the brain. One intuitive computational strategy for invariant face recognition is to separate information about facial shape from that about facial texture. Changes in head orientation or expression can produce changes in facial shape but leave unaltered the underlying texture map of the face (arising largely from physical features such as skin pigmentation, the shape and thickness of eyebrows, eyes, lips, and so on). An effective computational approach to decouple shape and texture information contained in a face is the “active appearance model,” a scheme for representing faces by projecting them onto two sets of axes, one describing the shape and one describing the shape-free appearance of a face (Cootes et al., 2001; Edwards et al., 1998). The decoupling between shape and texture parameters accomplished by the active appearance model approximately aligns with the needs of invariant face identification (though some shape-related features, e.g., inter-eye distance can vary depending on facial identity, and some appearance-related features, e.g., illumination, can vary for the same facial identity).

A recent study used facial images synthesized by an active appearance model to explore the coding scheme of AM face cells (Chang and Tsao, 2017). The study found that the active appearance model provides a remarkably simple account of AM activity: AM cells approximately encode linear combinations of axes of this model. However, this study left several issues unaddressed. First, the study used synthetic faces generated by the active appearance model rather than real faces. The code for real faces in the macaque brain may be different from that for synthetic faces. Furthermore, since the faces tested were directly controlled by parameters of the active appearance model, this may have given an unfair advantage to this model over other models for explaining face cell responses. Second, while the study compared the active appearance model to a few other models, it notably did not evaluate state-of-art deep networks trained on face recognition. Convolutional neural networks (CNNs) trained to perform face recognition now achieve close-to-human or even better performance (Parkhi et al., 2015; Taigman et al., 2014), naturally raising the question, how similar are the representations used by these artificial networks compared to those used by the primate face patch system?

Here, we set out to compare a large set of different models for face representation in terms of their power to explain neural responses from macaque face patch AM to pictures of real faces. The models tested include the original active appearance model used by Chang and Tsao (Chang and Tsao, 2017), referred to below as the “2D Morphable Model”, an Eigenface Model (Sirovich and Kirby, 1987; Turk and Pentland, 1991), a 3D Morphable Model (Blanz and Vetter, 1999; Paysan et al., 2009), several CNN models (Krizhevsky et al., 2012; Parkhi et al., 2015; Simonyan and Zisserman, 2015; Kubilius et al., 2018), a β variational autoencoder (Higgins et al., 2017), and a model implementing Hebbian learning on V1-like representations (Leibo et al., 2017).

## Results

We collected 2100 real faces from multiple online face databases, including the FERET face database (Phillips et al., 2000; Phillips et al., 1998b), CVL face database (Solina et al., 2003), MR2 face database (Strohminger et al., 2016), PEAL face database (Gao et al., 2008), AR face database (Martinez and Benavente, 1998), Chicago face database (Ma et al., 2015), and CelebA database (Yang et al., 2015) (Figure 1A). Responses of 159 face-selective cells in macaque face patch AM were recorded from two monkeys while presenting the facial images (Figure 1B). To find the optimal set of axes explaining neuronal responses, we extracted feature vectors from several different models, including 2D Morphable Model (Cootes et al., 2001), 3D Morphable Model (Blanz and Vetter, 1999), Eigenface Model (Sirovich and Kirby, 1987; Turk and Pentland, 1991), AlexNet (Krizhevsky et al., 2012), VGG-face (Parkhi et al., 2015), VGG-19 (Simonyan and Zisserman, 2015), CORnet (Kubilius et al., 2018), β-VAE (Higgins et al., 2017), and a model implementing Hebbian learning on V1-like representations (Leibo et al., 2017; Figure 1C). These models each parameterize faces using very different principles. The Eigenface model has the simplest form, consisting of principal components of pixel-level representations of facial images. The 2D and 3D Morphable Models are generative face models that convert a set of parameters into a facial image. AlexNet, VGGs, and CORnet are neural network models each trained on a different task: AlexNet, VGG-19, and the CORnets are all trained to classify images into 1000 non-face object categories. VGG-face is trained to identify 2,622 celebrities..The CORnet family includes three networks: CORnet-Z, CORnet-R, and CORnet-S. All three models have four areas that are identified with cortical areas V1, V2, V4, and IT. CORnet-Z has a purely feedforward structure, while CORnet-R and S contain recurrent connections within areas. β-VAE is a deep generative model that learns to faithfully reconstruct the input images, while being additionally regularized in a way that encourages individual network units to code for semantically meaningful variables. The Hebbian learning model is a biologically plausible model accounting for mirror-symmetric view tuning in face patch AL and view invariance in face patch AM. We chose these models because they are well known CNN models trained on object categorization (AlexNet, VGGs and CORnets), important computational models for face recognition (Eigenface, 2D Morphable Model, 3D Morphable Model, and Hebbian learning model), or a state-of-art neural network model for unsupervised disentangled representation learning (β-VAE). Note that the models being compared are quite different--the 2D/3D Morphable models are generative models for faces that do not possess any explicit identification with a specific stage of visual processing, while deep network models have an architecture in rough correspondence with the ventral visual stream (Fukushima, 1980).

**Figure 1.**
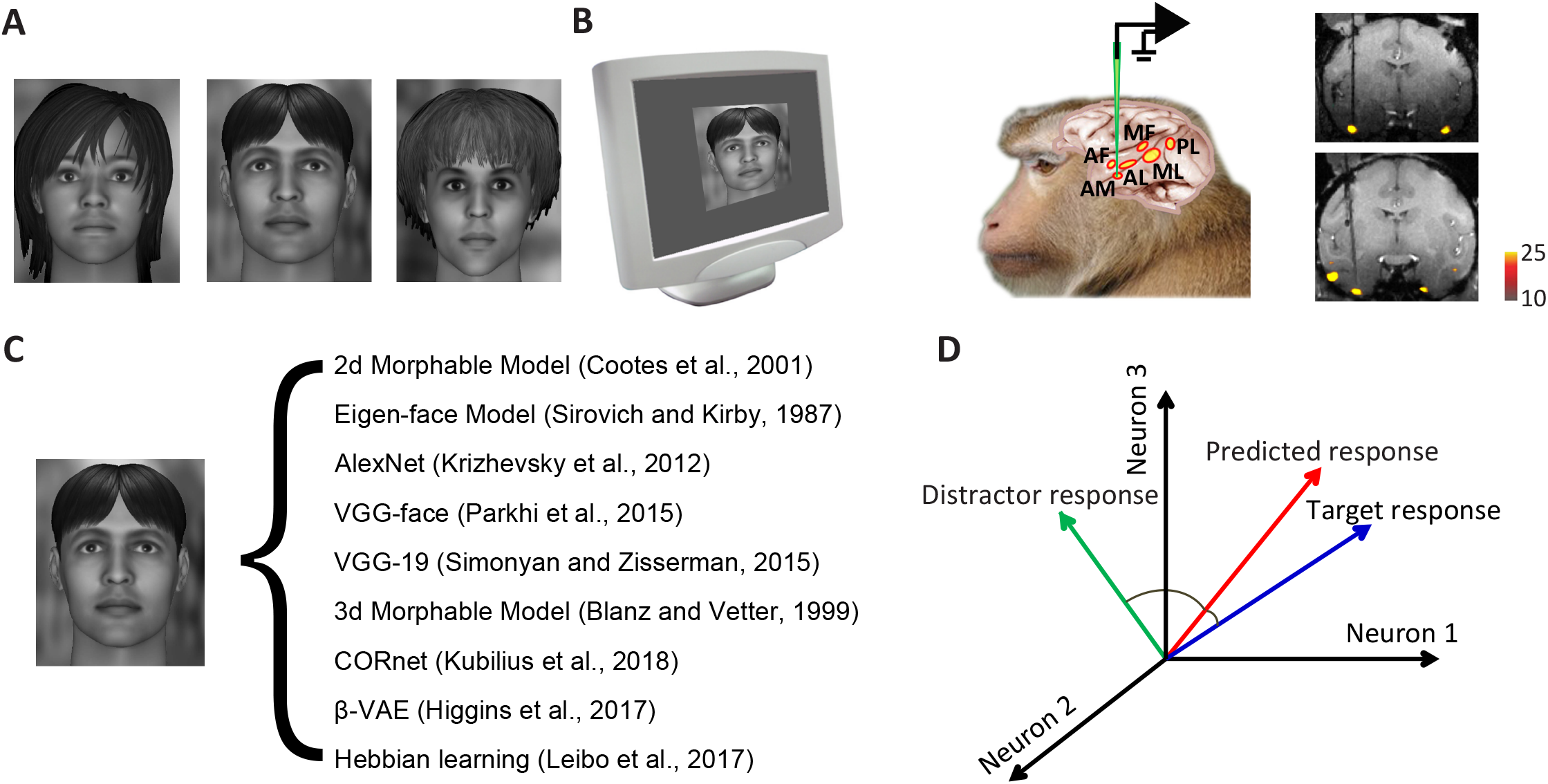
Stimulus and analysis paradigm. A, 2100 facial photos from multiple face databases were used in this experiment. Three examples are shown. [Note that the facial images shown here and in Figures 2–4 are synthetically generated faces that serve as stand-ins for the actual example faces in order to satisfy bioRxiv’s policy on the use of images of human faces; they are not from the database and were not shown to the monkeys.] B, Images were presented to the animal while recording from the most anterior face patch AM (anterior medial face patch). The electrode track targeting AM is shown in coronal MRI slices from two animals. C, Each facial image was analyzed using 9 different models. The same number of features were extracted from units of different models using principal component analysis (PCA) for comparison. D, Different models were compared with respect to how well they could predict neuronal responses to faces. A 10-fold cross-validation paradigm was employed for quantification: 2100 faces were evenly distributed into 10 groups. Responses of each neuron to 9 groups were fit by linear regression using features of a particular face model, and the responses of this neuron to the remaining 210 faces were predicted using the same linear transform. To quantify prediction accuracy, we compared predicted responses to individual faces in the space of population responses to either the actual response to that face or that to a distractor face. If the angle between predicted response and target response was smaller than that between predicted response and distractor response, the prediction was considered correct. All pairs of faces were used as both target and distractor and the proportion of correct predictions was computed.

To quantify how well each model can explain AM neuronal responses, for each model, we learned a linear mapping between features of that model and the neural population response vector. To avoid overfitting, we first reduced the dimensionality of each model by performing principal components analysis (PCA) on model responses to the 2100 faces, yielding N features for each face and each model. Then a 50-fold cross-validation paradigm was performed: responses of each neuron to 42*49=2058 faces were fit by linear regression using the N features, and then the responses of the neuron to the remaining 42 faces were predicted using the same linear transform. Besides measuring how much variance in neural responses could be explained by the linear transform for each individual neuron, we also took a different approach using the population response: we compared the predicted population response vector to each face with the actual population response vector to the face as well as the population response vector to a random distractor face (Figure 1D). If the angle between prediction and target was smaller than that between prediction and distractor, the prediction was considered correct.

To compare different models, we used the top 50 PCs of features from each model, as in a previous study (Chang and Tsao, 2017). We found the best model was one of the CORnet models, CORnet-Z, followed by the 2D Morphable Model (Figure 2A, good performance=high explained variance and low encoding error). Interestingly, we found that VGG-face, a deep network trained to identify individual faces, performed worse than the other models, while CORnet-Z, which was trained to classify 1000 classes of non-face objects, performed better than any other model. A confounding factor is that the images we used came from multiple face databases which have variable backgrounds. Some of the models may use background information more than other models for prediction. Hence performance differences between models could have been driven by representation of the background. In theory, the background should not be relevant to a model of face representation.

**Figure 2.**
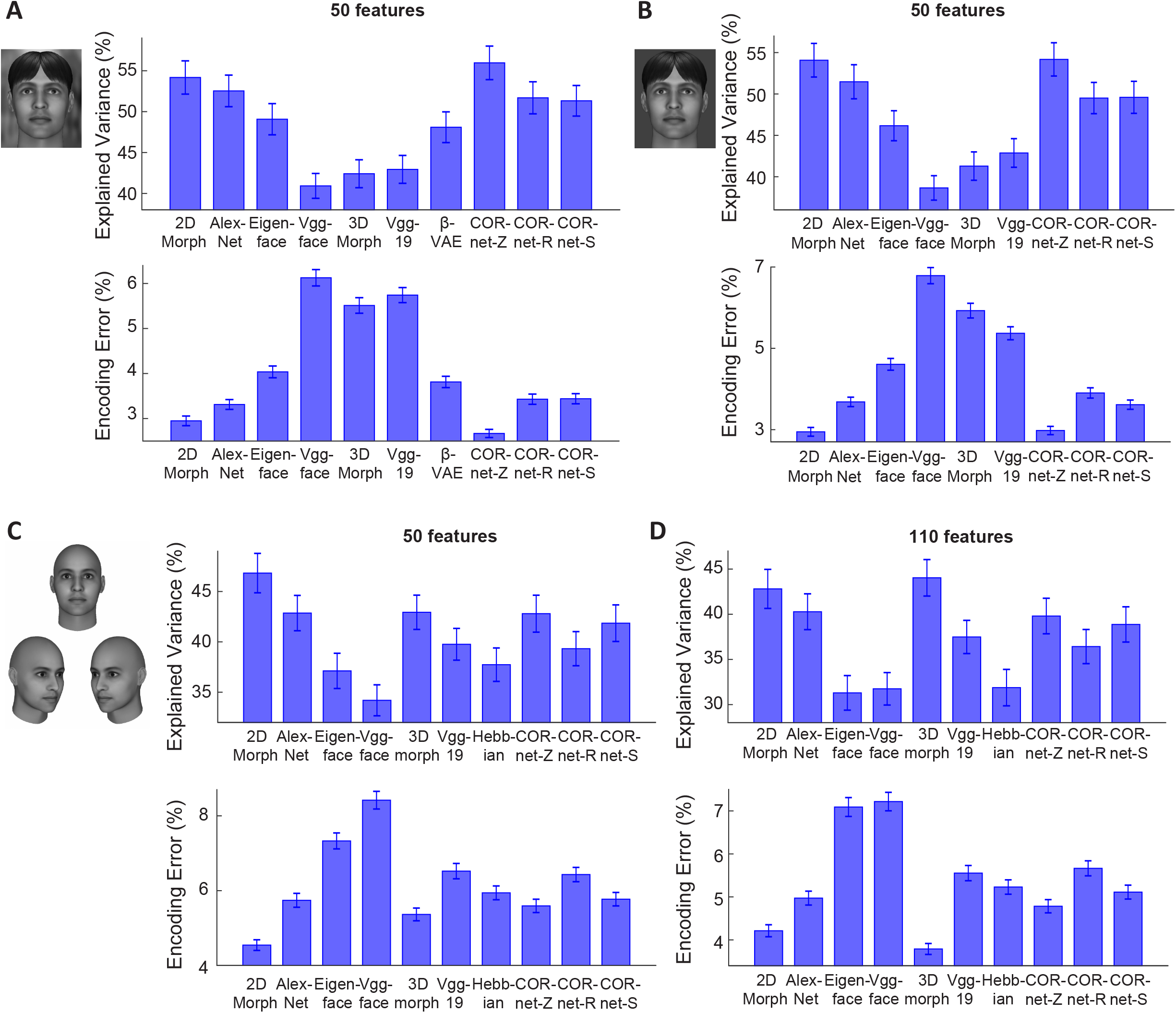
Comparing how well different models of face coding can explain AM neuronal responses to facial images. A, For each model, 50 features were extracted using PCA and used to predict responses of AM neurons. *Upper:* Explained variances are plotted for each model. For each neuron, explained variance was normalized by the noise ceiling of that neuron (see STAR Methods). Error-bars represent s.e.m. for 148 cells. CORnet-Z performed significantly better than the other models (p<0.001 in all cases except from the 2D Morphable Model, p<0.01 between CORnet-Z and the 2D Morphable Model, Wilcoxon signed-rank test), and the 2D Morphable Model performed significantly better than the remaining models (p<0.01). *Lower:* Encoding errors are plotted for each model. Error-bars represent s.e.m. for 2100 target faces (i.e., error was computed for each target face when comparing to 2099 distractors, and s.e.m was computed for the 2100 errors). CORnet-Z performed significantly better than the other models (p<0.001 in all cases except from the 2D Morphable Model, p<0.01 between CORnet-Z and the 2D Morphable Model, Wilcoxon signed-rank test), and the 2D Morphable Model performed significantly better than the remaining models (p<0.001). B, To remove differences between models arising from differential encoding of image background, face images with uniform background were presented to different models (see STAR Methods). CORnet-Z and 2D Morphable Model performed significantly better than the other models (p<0.001), with no significant difference between the two models (p=0.30 for explained variance; p=0.79 for encoding error). C, To create facial images without hair, each facial image in the database was fit using a 3D Morphable Model (left). [Note that the 3D morph fit shown here and in subsequent figures is a synthetically generated face that serves as a stand-in for the actual example 3D morph fit in order to satisfy bioRxiv’s policy on the use of images of human faces.] The fits were used as inputs to each model. For example, a new 2D Morphable Model was constructed by morphing the fitted images to an average shape. 50 features were extracted from each of the models using PCA for comparison. D, Same as C, but for 110 features. For 50 features, the 2D Morphable Model performed significantly better than the other models (p<0.001), while there was no significant difference between 3D Morphable Model and CORnetZ (p=0.19 for explained variance; p=0.21 for encoding error) or between 3D Morphable Model and AlexNet (p=0.56 for explained variance; p=0.06 for encoding error). For 110 features, the 3D Morphable Model outperformed all other models (p<0.01 between 2D Morphable Model and 3D Morphable Model for explained variance; p<0.001 in all other cases).

Thus we next extracted features from facial images without background (see STAR Methods), and repeated the comparison of different models. The ordering of model performance was largely preserved after background removal (Figure 2B). However, after background removal, the performance of the 2D Morphable Model was not significantly different from CORnet-Z (p=0.30 for explained variance; p=0.79 for encoding error); this is consistent with the fact that the 2D Morphable Model only accounts for intensity variations of faces, but not background.

In the above two cases, we found the performance of the 3D Morphable Model was much lower than the 2D Morphable Model. However, there is an important difference between the 2D and 3D Morphable Models: the latter does not fit hair-related features. To compensate for this difference, we further tested the models on hairless facial images derived from fits using the 3D Morphable Model (Figure 2C, left). We performed the analysis using either 50 PCs or 110 PCs (the dimension of the 3D Morphable Model). In both cases, the 3D Morphable Model outperformed VGG-face, VGG-19, CORnet-R, CORnet-S, Eigenface (Figure 2C). In the case with 110 PCs, it even outperformed AlexNet, CORnet-Z and the 2D Morphable Model (Figure 2D). For faces without hair, the 2D Morphable Model also performed significantly better than all of the neural network models. As an alternative strategy to removing hair, a mask was created by the 3D morphable model fit, and the original image was cropped using that mask, thus ensuring that all non-hair features were left unchanged. The results were largely consistent (Figure S1).

Furthermore, the use of the facial images generated by the 3D Morphable Model allowed us to test a Hebbian learning model recently proposed to account for face patch responses (Leibo et al., 2017). This model posits that the weights of face cells, learned through Hebbian learning, converge to the top PCs of the neuron’s past inputs, and these inputs should generically constitute short movies of faces rotating in depth. The 3D Morphable Model allowed us to readily synthesize a set of facial images at multiple views and thus test the Hebbian model (see STAR Methods). For explained variance, the Hebbian model performed better than VGG-face (p<0.001 for 50 PCs, but p=0.82 for 110 PCs), comparably to Eigenface (p=0.32 for 50 PCs, p=0.35 for 110 PCs), and worse than the other models (Figure 2C and D, upper panels). For encoding error, the Hebbian model performed better than the VGGs, CORnet-R and Eigenface models, comparably to CORnet-S (p=0.11 for 50 PCs and p=0.37 for 110 PCs) and AlexNet (p=0.16 for 50PCs, p=0.06 for 110 PCs), and worse than the 2D Morphable Model, 3D Morphable Model, and CORnet-Z (Figure 2C, D).

Finally, we performed a more detailed comparison between the 2D Morphable Model and another generative model, β-VAE, whose latent units are encouraged to encode semantically meaningful variables, otherwise known as disentangled variables (Higgins et al., 2017). β-VAE contains an encoder that transforms the image input into a vector of disentangled latent variables and a decoder that transforms the vector back into an image. The encoder is implemented with a convolutional neural network, similar to other network models. A recent analysis of the same data set as in the present paper found that a subset of single AM cells have selectivity remarkably matched to that of single β-VAE latents, suggesting that AM and β-VAE have converged, at least partially, upon the same set of parameters for describing faces (Higgins et al., 2020). In particular, the single-neuron alignment between AM and β-VAE was better than that for any other model including the 2D morphable model. Thus we wanted to address in detail how β-VAE compares to the 2D Morphable Model by the metric of neural population encoding (Figure 2A). Due to variations in training parameters, 400 different β-VAE models were trained, each with 50 latent units. For the comparison in Figure 2A, we chose the β-VAE with the least encoding error. We further compared encoding performance of all 400 β-VAE models to that of the 2D morphable model (Figure S2A). Close inspection revealed that some of the latent units have much smaller variance than other units in response to 2100 faces, thus we removed those units with variance <0.01. We found for both β-VAE and 2D Morphable Model explained variances increased and encoding errors decreased when the dimensionality increased, as one might expect, since more dimensions are available for capturing the information present in the neural responses, and the 2D Morphable Model performed better than β-VAE models at matched feature dimensions (Figure S2A and C, p<0.05 in all cases, p<0.01 except from dimension=16 for encoding error). However, among the 400 β-VAEs reported in Figure S2, many models did not learn a well disentangled representation, hence failing at one of the optimisation objectives. Therefore, we also compared the 2D Morphable Model to a subset of β-VAEs where the units were well disentangled (based on the unsupervised disentangled ranking (UDR) score for each β-VAE, see STAR Methods), and found that the encoding performance difference was not significant at low dimensions, but the 2D Morphable Model performed better for feature dimensions=6, 10 and 12 (Figure S2A and C, *inset*). Thus we conclude that by the metric of explained variance and neural encoding performance, the 2D Morphable Model outperforms other generative models, including both β-VAE and Eigenface models, and performs similarly to disentangled β-VAEs at lower dimensions.

Next, we asked the complementary question: how well could we predict the model features by linear combinations of neural responses of face cells? The same procedure as the encoding analysis was followed, except that the respective roles played by model features and neural responses were reversed. The results for this decoding analysis were largely consistent with encoding analysis (Figure 3). For the original images and images without the background, the 2D Morphable Model performed better than all other models except CORnet-Z (p=0.08 for original images, p=0.03 after background removal with 2D Morphable Model performing slightly better). For images generated by the 3D Morphable model, the 2D Morphable Model outperformed all other models except the 3D Morphable model (p=0.42 for 50-d model, p<0.001 for 110-d model). Comparing the 2D Morphable Model with β-VAEs at matched dimensions, we found that the two models were comparable at dimensions≤16 (Figure S2E), but the 2D Morphable Model performed better at higher dimensions. The fact that the performance of the two models were more comparable in both decoding and encoding at low dimensions (Figures S2A, C and E) suggests that with certain training parameters, β-VAE was efficient at extracting a small number of meaningful features, but the number of disentangled features discovered in this way may not be sufficient to achieve a good performance in decoding/encoding. To better illustrate the relationship between feature dimensionality, encoding/decoding errors and disentanglement, we quantified the quality of disentanglement achieved by β-VAE models by the UDR score as before. We found positive correlations between the encoding/decoding errors and UDR (Figure S2D and F), and negative correlations between the explained variance, the number of informative features and UDR (Figure S2B and G). These results support the idea that the objective of disentanglement encourages the model to converge to a small number of informative features, at the expense of the overall explanatory power of the face code in the brain.

**Figure 3.**
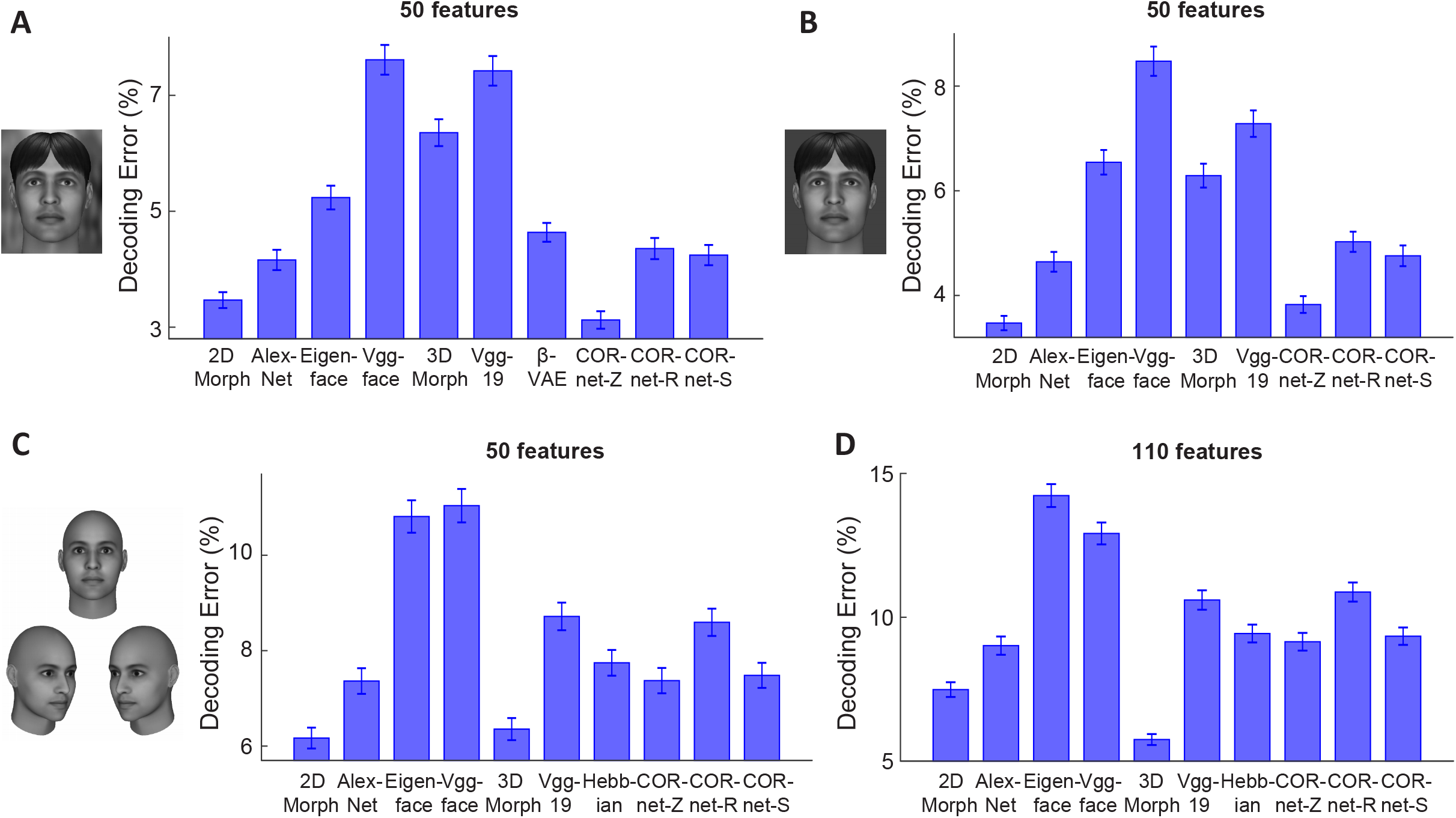
Comparing how well AM neuronal responses to facial images can explain different models of face coding. A, For each model, 50 features were extracted using PCA and responses of AM neurons were used to predict the model features. Decoding errors are plotted for each model. Error-bars represent s.e.m for 2100 target faces (i.e., error was computed for each target face when comparing to 2099 distractors, and s.e.m was computed for the 2100 errors). CORnet-Z performed significantly better than the other models (p<0.001) except from the 2D Morphable Model (p=0.08, Wilcoxon signed-rank test), and the 2D Morphable Model performed significantly better than the remaining models (p<0.01). B, To remove differences between models arising from differential encoding of image background, face images with uniform background were presented to different models (see STAR Methods). CORnet-Z and 2D Morphable Model performed significantly better than the other models (p<0.001), with only a small difference between the two models (p=0.03). C, To create facial images without hair, each facial image in the database was fit using a 3D Morphable Model (left). The fits were used as inputs to each model. For example, a new 2-D Morphable Model was constructed by morphing the fitted images to an average shape. 50 features were extracted from each of the models using PCA for comparison. D, Same as C, but for 110 features. For 50 features, the 2D Morphable Model and 3D Morphable model performed significantly better than the other models (p<0.001), with no significant difference between the two models (p=0.42). For 110 features, the 3D Morphable Model outperformed all other models (p<0.001).

So far, for our deep network comparisons, we have focused on comparing how well units in the penultimate layer of each deep network explain AM neural responses (Yamins et al., 2014; Cadieu et al., 2014). We also examined how well each individual layer of AlexNet and CORnets explained neural responses (Figure S3). Surprisingly, we found intermediate layers of those models performed the best (layer L2 of AlexNet and the V4 layer of CORnet-Z) (Figure S3A-D). However, when we examined the relationship between different layers of AlexNet and AM responses using a second stimulus set containing facial images at 8 head orientations, we found that AlexNet L2 lost its advantage, with L5/6 performing the best (Figure S3E and F). Given that AlexNet L2 cannot explain neural responses to a stimulus set with different head orientations, it is clearly not a viable candidate for explaining face patch activity.

Overall, our results suggest that linear combinations of features of the 2D Morphable Model were closely related to the responses of face cells, achieving encoding and decoding performance comparable to even the best neural network models developed recently (Figures 2 and 3), while at the same time using a simple and transparent representation that does not involve hundreds of thousands of “black box” parameters. This result also extends our previous finding from synthetic faces to real faces (Chang and Tsao, 2017). The similar performance of several of the models to the 2D Morphable Model (e.g., AlexNet and CORnet-Z in Figure 2A and 3A) raises the question whether these other models are simply linear transformations of the 2D Morphable Model, or whether they provide “additional” features that could help explain neural responses.

To address this question, we concatenated features from two different models, and asked how much more variance in neural responses could be explained by two models in comparison to a single model (see STAR Methods) (Figure 4A). This quantified the neural variance uniquely explained by a model (i.e., unique explained variance by model A = explained variance by models A and B - explained variance by model B). Overall, we found unique variances explained by various models compared to the 2D Morphable Model to be much smaller than that by full models (compare Figure 4A to the upper panel of Figure 2B), suggesting other models provide limited “additional” features to explain neural responses. For facial images without background, AlexNet accounted for the most unique variance across all models, followed by CORnet-Z, CORnet-R, CORnet-S, VGG-19, VGG-face, 3D Morphable Model and Eigenface (Figure 4A). For hair-free reconstructions, CORnet-Z accounted for the most unique variance across all models, followed by CORnet-S, AlexNet, CORnet-R, VGG-19, 3D Morphable Model, VGG-face and Eigenface (Figure 4C). Furthermore, we asked the opposite question: what happens when the 2D Morphable Model is combined with other models? Again, we found unique variances explained by the 2D Morphable Model to be much smaller than the full 2D Morphable Model, indicating that the 2D Morphable Model features significantly overlap those of other models. However, a significant extent of non-overlap was also found (Figure 4B). For facial images without background, the most unique variance was explained by comparing the 2D Morphable Model to VGG-face, followed by VGG-19, 3D Morphable Model, CORnet-R, CORnet-S, Eigenface, AlexNet and CORnet-Z (Figure 4B). For hair-free reconstructions, the most unique variance was explained by comparing the 2D Morphable Model to VGG-face, followed by CORnet-R, Eigenface, VGG-19, CORnet-S, AlexNet, CORnet-Z and 3D Morphable Model (Figure 4D). Finally, we asked how much unique variance each layer of AlexNet explained compared to the 2D Morphable Model by repeating the analysis of Figure 4A on individual layers of AlexNet. We found that the amount of unique variance was similar across layers L2 to L6 (Figure S3G). However, the unique variance explained by the 2D Morphable Model compared to each of the layers of AlexNet was not equal across layers, but was minimal for layer L2 (Figure S3H). This may partially explain why AlexNet layer L2 could explain more variance than other layers for the 2100 frontal face set (Figure S3A-D).

**Figure 4.**
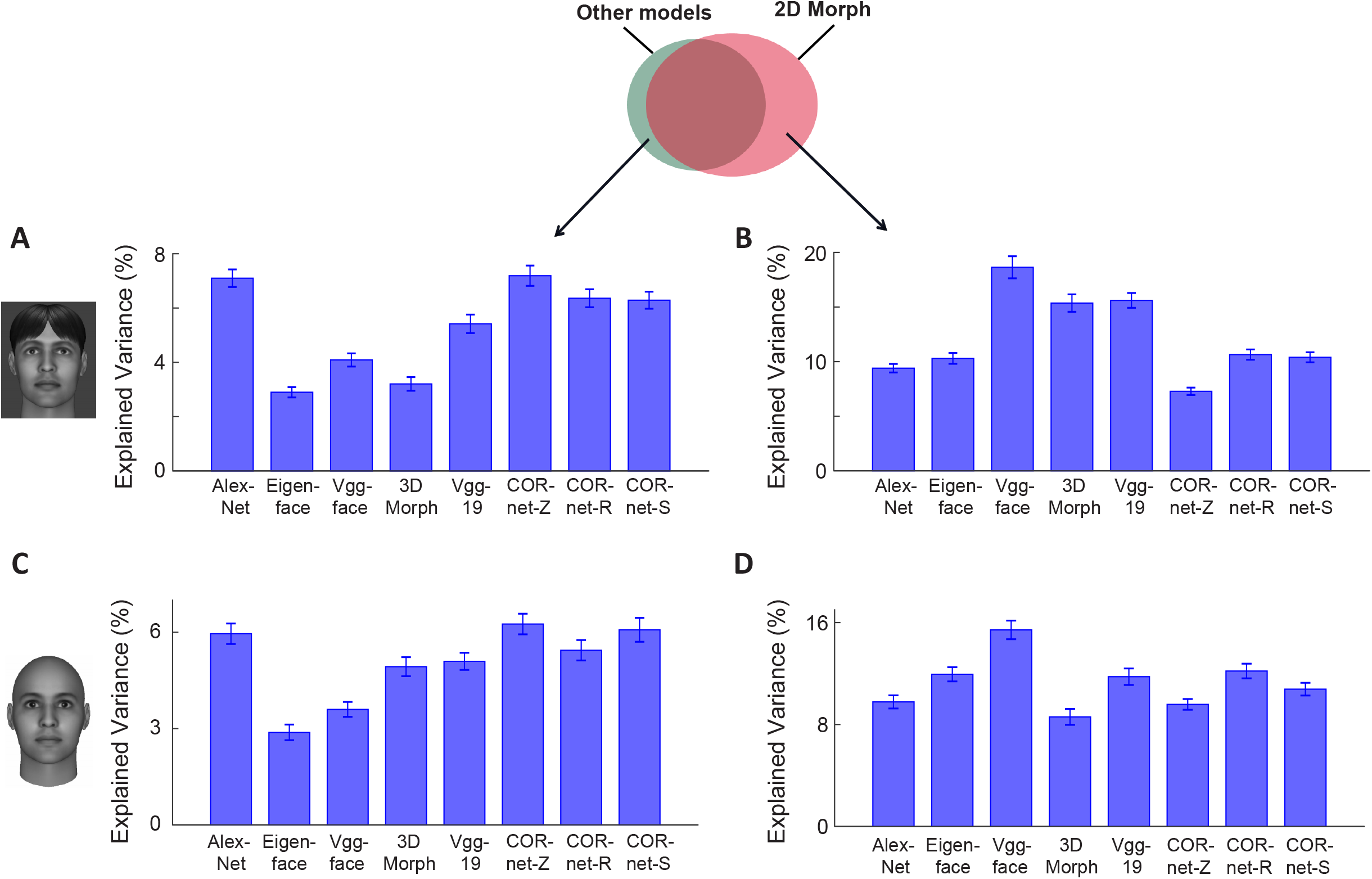
Measuring neural variance uniquely explained by the 2D Morphable Model and other models. Fifty model features from the 2D Morphable Model and a different model were concatenated, and neural responses were fit using all 100 features as regressors. The explained variance was then subtracted by the contribution from each individual model, to quantify the efficacy of the non-overlapping components of the two models in predicting neural responses. A, Percentage of neural variance uniquely explained by various models compared to the 2D Morphable Model, for images after background removal (cf. Figure 2B). B, Percentage of neural variance uniquely explained by the 2D Morphable Model compared to other models. C and D, same as A and B, but for images fit by the 3D Morphable Model (cf. Figure 2C). Error-bars represent s.e.m. for 148 cells.

In the above analysis, models were compared with respect to the amount of variance they could explain in the neural responses. We also asked how well each model could explain features of other models through linear regression, independent of neural responses. To address this, we repeated the analysis of Figure 4, but instead of using real neurons, we directly quantified how much variance of features in one model could be explained by features of the 2D Morphable Model (Figure S4A and B). Features of the 2D Morphable Model only partially explain features of the other models (Figure S4A, blue bar), suggesting the models do provide additional features beyond those of the 2D Morphable Model. However, not all of these features help to explain neural responses. For example, the Eigenface model contains a sizable component unexplained by the 2D Morphable Model (Figure S4A), but the amount of neural variance explained by the adding the Eigenface model to the 2D Morphable Model was only marginal (Figure 4A). We also asked how well each of the other models explain 2D Morphable features (Figure S4B). The most variance was explained by Eigenface, while the least variance was explained by Vgg-face. Finally, to compare all model pairs on equal footing, Figure S4C plots the amount of variance in each 50-feature face model explained by each of the other models. Interestingly, the 2D Morphable Model explained as much variance in CORnet-Z features as AlexNet. This shows that for the subspace of faces, an explicit generative model of face representation, the 2D Morphable Model, can do as good a job at explaining CORnet-Z features as a deep network explicitly built with similar architectural principles and training procedure (AlexNet).

Overall, the 2D Morphable Model is composed of two components: a shape component defined by positions of facial landmarks, and a shape-free appearance (or texture) component defined by the intensity distribution of the facial image after shape normalization. Similarity between neural network models (esp. CORnets) and the 2D Morphable Model indicates the networks may implement similar computation to the 2D Morphable Model, by first localizing the landmarks, then morphing the face to remove shape-related information. We investigated how these two components were related to different stages of neural network models (Figure S4D-G). We found that one intermediate layer, layer V4, in CORnets best explains the shape component, consistent with the interpretation that the intermediate stage may perform the computation of finding landmarks. When examining the shape-free appearance component, however, we were surprised to find the input to the networks (i.e., pixel intensities of the images) performed the best across all layers of CORnets (Figure S4D). Inspection of the spatial profile of decoding suggests the input layer well captured global features such as the intensity of the skin (Figure S4F), rather than local features, such as the eyes and the mouth, which is quite different from the neural data (Figure S4G, left). Indeed, the correlation between decoding maps reveals an increase in similarity with neurons across different layers of CORnets (Figure S4G, right). This analysis suggests different stages of CORnets are performing something similar to different stages of the 2D Morphable Model, at least for the purpose of explaining neural responses.

Finally, we wanted to gain some insight into why AlexNet outperforms VGG-face in explaining neural responses, demonstrated by both the encoding and the decoding analyses (Figures 2, 3). This is surprising since AlexNet is not trained to classify any face images (albeit some images within the different training classes do contain faces), while Vgg-face is trained exclusively to identify face images. We started by computing similarity matrices (Kriegeskorte et al., 2008) for neural population responses from face patch AM (Figure 5A1). Here each entry of the matrix represents the similarity between a pair of faces, quantified as correlation between population responses to the face pair. The same analysis was repeated with the top 50 PCs of deep features from AlexNet or VGG-face (Figure 5A2, A3). There is a clear difference between the similarity matrix for VGG face compared to those for AM and Alexnet features. Similarity matrices for both AM and AlexNet features show a dark cross with a bright center, but this is not the case for VGG face. We computed the difference between the Vgg-face and AlexNet matrices (Figure 5A4), and then shuffled the rows and columns according to the first principle component of the difference matrix (Figure 5A5). After sorting, positive entries tended to be located at the bottom-left and the upper-right corner (square outlines in Figure 5A5): here a positive difference indicates the two faces are more similar under features of VGG-face than AlexNet; therefore the faces at the opposite ends of PC1 are more likely to be confused by VGG-face. What do the faces at the two extremities look like? To examine the difference, we picked the first 100 faces and last 100 faces along the direction of PC1, and divided them into 20 groups of 10 faces. An average face after shape normalization was generated for each group (Figure 5B). We see an interesting difference: The first 10 groups of faces show inhomogeneous illumination--some parts of faces, such as cheeks and hair, are brighter than other parts of the face, such as the mouth, while the last 10 groups of faces appear more homogeneously illuminated.

**Figure 5.**
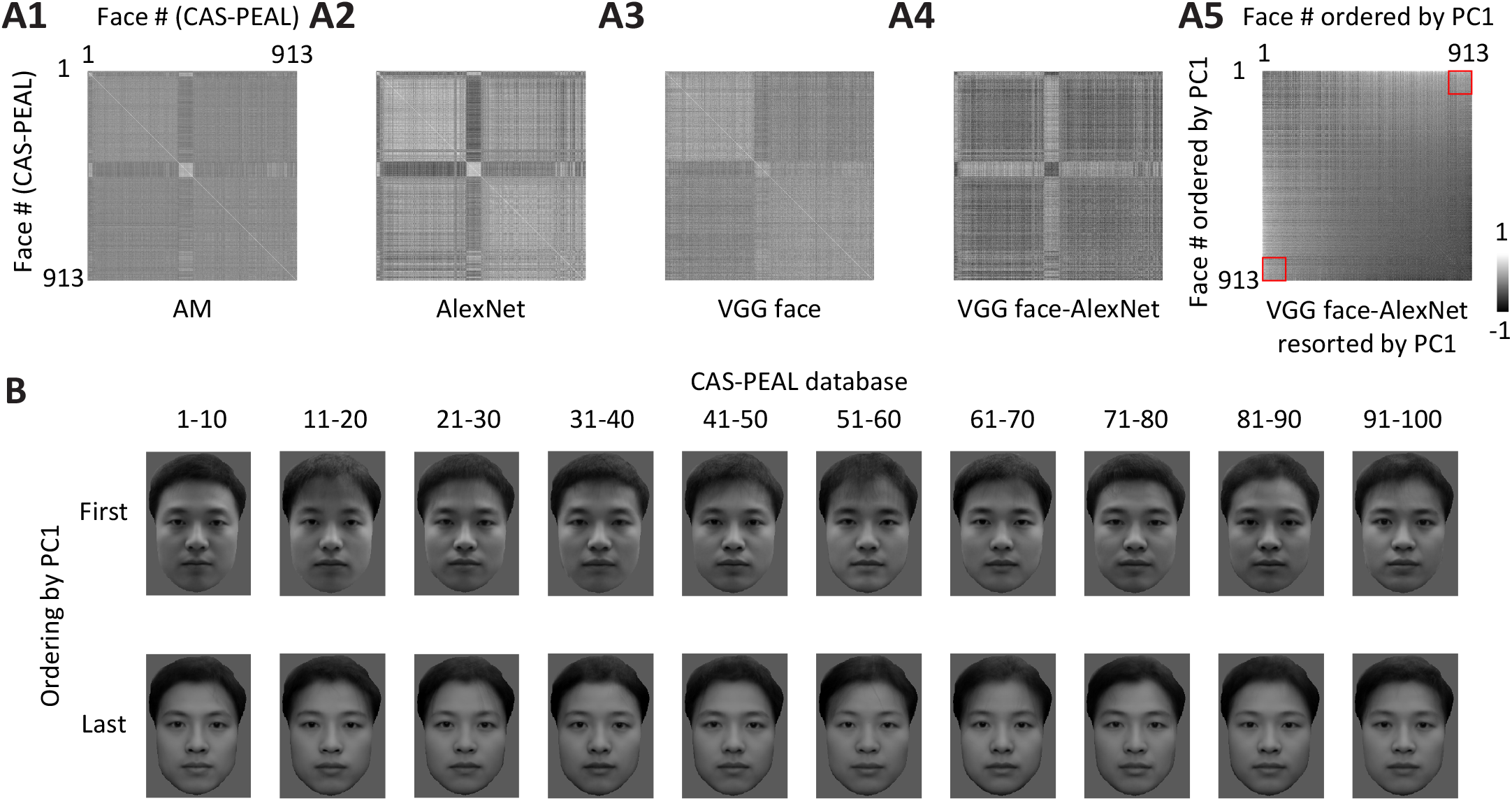
Vgg-face features and AlexNet features show a marked difference in coding illumination levels. A, Similarity matrices were computed for 913 faces from CAS-PEAL database using AM population responses (A1) and features of two network models, AlexNet (A2) and Vgg-face (A3). Each entry indicates the correlation between representations of two faces. The difference between the two matrices derived from the network models was computed (A4). Rows and columns of the differential matrix were shuffled according to the first principal component of the difference matrix (A5). The red squares outline face pairs taken from the first and last 100 faces: these face pairs showed a significantly higher representational similarity by Vgg-face compared to AlexNet. B, First 100 faces and last 100 faces along the direction of PC1 were divided into 20 groups of 10 faces. An average face after shape normalization was generated for each group. [Note that images shown here are not actual faces of any individuals, but the average images of 10 faces after being morphed to the average shape, using the same algorithm as the 2D Morphable Model.]

In the analyses of Figure 5, we used a database, CAS-PEAL, which contains only Chinese faces. Is this observation unique to Chinese faces? We repeated the same analysis for 748 Caucasian faces. Similar to CAS-PEAL faces, we found that the face groups eliciting a much more similar representation by Vgg-face compared to AlexNet consisted of faces with unbalanced versus homogeneous illumination (Figure S5). In sum, we found that VGG-face is much less sensitive to illumination differences than both AM cells as well as AlexNet, and this likely contributes to the inferior ability of Vgg-face to predict AM responses compared to AlexNet.

## Discussion

Face processing has been a subject of intense research effort in both visual neuroscience and computer vision, naturally raising the question, what, if any, computer vision model of face representation best matches that used by the primate brain. A recent paper found the 2D Morphable Model, a classic model of face representation from computer vision, could explain neural activity in face patches remarkably well (Chang and Tsao, 2017). At the same time, a number of groups have found that activity in deep layers of convolutional neural networks can explain significant variance of neural responses in ventral temporal cortex (Yamins et al., 2014; Kalfas et al., 2017; Yildirim et al., 2020; Schrimpf et al., 2018; Raman and Hosoya, 2020). Here, we extend those results by comparing the efficacy of a large number of different computational models of face representation to account for neural activity in face patch AM. We were especially interested in how the 2D morphable model, a simple and explicit graphical model, would compare to Vgg-face, a black box deep neural network dedicated to face recognition containing hundreds of thousands of parameters and trained on nearly a million (982,803) facial images. Our findings suggest that the 2D Morphable Model is better than most other models in explaining the neuronal representation of real faces including Vgg-face. For faces without background, the 2D Morphable Model allowed better linear coding of neural responses by model features than every model except CORnet-Z, whose performance it matched, with differences depending on presence of background and hair (specifically, the 2D Morphable Model performed worse than CORnet-Z for faces with both hair and backgrounds, comparably to CORnet-Z for faces with no backgrounds, and better than CORnet-Z for faces without hair reconstructed by 3D Morphable Model). This is surprising, since the 2D Morphable Model is one of the oldest models for face representation (next to the Eigenface model). Furthermore, the 2D Morphable Model matched AlexNet in its ability to explain CORnet-Z features (Figure S4C, row 7). Since AlexNet is a deep network whose architectural principles and training procedure CORnet-Z explicitly emulates, these results suggest that the face subspace portion of the representation learned by CORnet-Z may be interpreted in simpler and more explicit terms, as a shape appearance model. The results provide an important counter-example to the increasingly popular view that only distributed representations learned by multi-layer networks can well explain IT activity (Kietzmann et al., 2018; Lillicrap et al., 2020). Why a network trained on object classification should learn an approximation to a generative model of faces is an interesting question for future research.

A recent theoretical study found that a deep neural network, when trained on certain tasks, gradually abandons information about the input unrelated to the task in its deep layers (Tishby and Zaslavsky, 2017). Thus artificial neural networks are unlikely to be fully identical to the brain, since the tasks both systems are trained on are unlikely to be identical. It makes sense that VGG-face is not able to distinguish illumination, since the identity of an individual does not depend on illumination. Why does AlexNet still contain information about illumination? It is possible this occurs because AlexNet has not been trained specifically on face identification, and illumination-related features are useful for more general object classification tasks (e.g., distinguishing a concave hole from a convex bump (Ramachandran, 1988)). In contrast, it appears that VGG-face is so specialized that any information unrelated to identity is filtered out in the end. The representation of illumination in AlexNet and its superior performance in predicting neural is in line with the observation that our recognition of unfamiliar faces is susceptible to changes in lighting conditions (Young and Burton, 2018).

The fact that object-general Cornet-Z model largely did as well as the face-specific 2DMorphable model in explaining face cell responses, much better than VGG-face, suggests that macaque face cells may not be entirely “domain-specific,” but have a more “generalist” function than previously understood. Within the face domain, face cells are not over-specialized for facial identification, but rather may provide an array of high-level information related to different aspects of faces in the visual field. Because the 2D Morphable Model retains all information needed to reconstruct a face, it should be able to perform well on any face-related task.

The finding that CORnet-Z was the best model among CORnet-Z, -R, and -S for encoding face cell responses was somewhat surprising, as CORnet-Z is the simplest model, with a purely feed-forward structure, while CORnet-R and CORnet-S are recurrent models (Kubilius et al., 2018). This could be a result of stimulus selection: the recurrent layers could help establish invariance in complex/difficult situations, but may not be an advantage in our case, since our stimuli are well aligned. This suggests the results of model comparison depend on the stimuli being used in the study. This also indicates the relatively narrow range of our stimulus space (mostly single faces with simple background) may limit the generality of our results. Further studies will require extending the comparison to more complex situations.

Overall, our analyses comparing a large number of models in terms of their ability to explain responses of cells in face patch AM show that a simple and explicit generative face model, the 2D Morphable Model, performs surprisingly well—rivaling or surpassing the deep network classifiers considered. This result supports the hypothesis that deep networks may be generally understood as inverting generative models (Lin and Tegmark, 2016; Ho et al., 2018). Several other lines of research investigating the representation of non-categorical variables in IT neurons have also converged on the same idea (Hong et al., 2016; Nishio et al., 2014). This raises the possibility the generative models, such as 2D Morphable Model, could help us reveal the mechanisms underlying face recognition in both the primate brain and the “black box” neural networks. Furthermore, the extremely poor performance of a deep network trained for face recognition in explaining face cell responses may give insight into the constraints shaping face patch development, raising the possibility that face patches may be optimized not for face recognition per se but instead for face reconstruction supporting arbitrary face-related behaviors.

## STAR Methods

### KEY RESOURCES TABLE

| REAGENT or RESOURCE | SOURCE | IDENTIFIER |
| --- | --- | --- |
| Experimental Models: Organisms/Strains |  |  |
| Rhesus macaques (*Macaca mulatta*) | UC Davis primate research center | N/A |
| Software and Algorithms |  |  |
| MATLAB | MathWorks | http://mathworks.com/ |
| MatConvNet | VLFeat | http://www.vlfeat.org/matconvnet |
| Basel face model | Gravis Research Group, University of Basel | https://github.com/unibas-gravis/basel-face-pipeline |
| Menpo | Intelligent Behaviour Understanding Group, Imperial College London | http://www.menpo.org |
| Other |  |  |
| Tungsten Microelectrode | FHC | Lot #:221355 |

## Resource Availability

### Lead Contact

Further information and requests for reagents and resources may be directed to Dr. Doris Tsao.

## Data and Software Availability

All data are available upon reasonable request to the Lead Contact.

## Experimental Model Details

Two male rhesus macaques (Macaca mulatta) of 7-10 years old were used in this study. Both animals were pair-housed and kept on a 14 hr/10hr light/dark cycle. All procedures conformed to local and US National Institutes of Health guidelines, including the US National Institutes of Health Guide for Care and Use of Laboratory Animals. All experiments were performed with the approval of the Caltech Institutional Animal Care and Use Committee (IACUC).

### Face Patch Localization

Two male rhesus macaques were trained to maintain fixation on a small spot for juice reward. Monkeys were scanned in a 3T TIM (Siemens, Munich, Germany) magnet while passively viewing images on a screen. Feraheme contrast agent was injected to improve signal/noise ratio. Face patches were determined by identifying regions responding significantly more to faces than to bodies, fruits, gadgets, hands, and scrambled patterns, and were confirmed across multiple independent scan sessions. Additional details are available in previous publications (Freiwald and Tsao, 2010; Ohayon et al., 2012; Tsao et al., 2006).

### Single-unit Recording

Tungsten electrodes (18–20 Mohm at 1 kHz, FHC) were back loaded into plastic guide tubes. Guide tubes length was set to reach approximately 3–5 mm below the dura surface. The electrode was advanced slowly with a manual advancer (Narishige Scientific Instrument, Tokyo, Japan). Neural signals were amplified and extracellular action potentials were isolated using the box method in an on-line spike sorting system (Plexon, Dallas, TX, USA). Spikes were sampled at 40 kHz. All spike data were re-sorted with offline spike sorting clustering algorithms (Plexon). Only well-isolated units were considered for further analysis.

### Behavioral Task and Visual Stimuli

Monkeys were head fixed and passively viewed the screen in a dark room. Stimuli were presented on a CRT monitor (DELL P1130). The intensity of the screen was measured using a colorimeter (PR650, Photo Research) and linearized for visual stimulation. Screen size covered 27.7*36.9 visual degrees and stimulus size spanned 5.7 degrees. The fixation spot size was 0.2 degrees in diameter and the fixation window was a square with the diameter of 2.5 degrees. Images were presented in random order using custom software. Eye position was monitored using an infrared eye tracking system (ISCAN). Juice reward was delivered every 2–4 s if fixation was properly maintained. For visual stimulation, all images were presented for 150 ms interleaved by 180 ms of a gray screen. Each image was presented 3–5 times to obtain reliable firing rate statistics. In this study, two different stimulus sets were used:

a. A set of 16 real face images, and 80 images of objects from nonface categories (fruits, bodies, gadgets, hands, and scrambled images) (Freiwald and Tsao, 2010; Ohayon et al., 2012; Tsao et al., 2006).
b. A set of 2100 images of real faces from multiple face databases, FERET face database (Phillips et al., 2000; Phillips et al., 1998b), CVL face database (Solina et al., 2003), MR2 face database (Strohminger et al., 2016), PEAL face database (Gao et al., 2008), AR face database (Martinez and Benavente, 1998), Chicago face database (Ma et al., 2015) and CelebA database (Yang et al., 2015). 17 online photos of celebrities were also included.
c. A set of 220 real face images, with 28 identities at 6-8 head orientations (Freiwald and Tsao, 2010).

### Quantification and Statistical Analysis

#### Selection of face selective cells

To quantify the face selectivity of individual cells, we defined a face-selectivity index as:

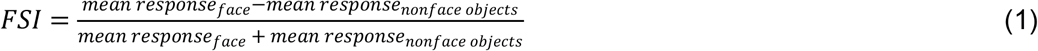

The number of spikes in a time window of 50-350 ms after stimulus onset was counted for each stimulus. Units with high face selectivity (FSI > 0.33) were selected for further recordings.

#### Extraction of facial feature from images

Each facial image was fed into the following models to extract corresponding features:

##### 1) 2D Morphable Model

This is the same model as used in our previous paper (Chang and Tsao, 2017) and feature extraction followed the procedure of previous papers on active appearance modeling (Cootes et al., 2001; Edwards et al., 1998). First, a set of 80 landmarks were labeled on each of the 2100 facial images. Out of the 80 landmarks, 68 were automatically labeled using an online package (“menpo”, http://www.menpo.org) and the remaining 12 were manually labeled. The positions of landmarks were normalized for mean and variance for each of the 2100 faces, and an average shape template was calculated. Then each face was smoothly warped so that the landmarks matched this shape template, using a technique based on spline interpolation (Bookstein, 1989). This warped image was then normalized for mean and variance and reshaped to a 1-d vector. Principal component analysis was carried out on positions of landmarks and vectors of shape-free intensity independently. Equal numbers of shape PCs and shape-free appearance PCs were extracted to compare with other models (25 shape/25 appearance PCs vs. 50 features of other models; 55 shape/55 appearance PCs vs. 110 features of other models). This model was also used to generate images without background used in Figure 2B. In this case, we first morphed all 2100 faces to the shape template, defined a mask for the standard shape to remove the background, and then morphed the masked facial image back to the original shape.

##### 2) 3D Morphable Model

We built a grayscale variant of the Basel Face Model (Paysan et al., 2009) from the original 200 face scans. The ill-posed 3D reconstruction from a 2D image was solved using (Schonborn et al., 2017) and the publicly available code from (Gerig et al., 2018). The first 50 principal components for the shape and color model respectively were adapted during the model adaptation process. The sampling-based method was initialized with the same landmarks as provided to the 2D Morphable Model. The pose was fixed to a frontal pose and the spherical harmonic illumination parameters were estimated robustly using (Egger et al., 2018) and the average illumination condition was fixed for the whole dataset. Note that the full complexity and flexibility of the 3DMM is not explored when analyzing frontal images only. Besides the model adaptation novel views where generated using the standard 3DMM pipeline by changing the head orientation and camera parameters. The images with varying head orientations were used to construct the Hebbian learning model (see below).

##### 3) Eigenface model

PCA was performed on the original image intensities of 2100 faces and top 50/110PCs were extracted to compare with other models.

##### 4) Pre-trained neural network models

We loaded 2100 facial images into the following pre-trained neural networks: (1) a MATLAB implementation of AlexNet: This network contains 8 layers: 5 convolutional layers and 3 fully connected layers, and has been pre-trained to identify a thousand classes of non-face objects. (2) a MATLAB implementation of Vgg-face neural network (Parkhi et al., 2015). This network contains 16 layers: 13 convolutional layers+3 fully connected layers, and has been pre-trained to recognize faces of 2622 identities. (3) a MATLAB implementation of Vgg-19 neural network (Simonyan and Zisserman, 2015). This network contains 19 layers: 16 convolutional layers and 3 fully connected layers, and has been pre-trained to identify a thousand objects. (4) a PyTorch implementation of CORnet (Kubilius et al., 2018). The CORnet family includes three networks: CORnet-Z, CORnet-R, and CORnet-S. All three models have four areas that are identified with cortical areas V1, V2, V4, and IT. CORnet-Z is the simplest model of the three, involving only feedforward connections, CORnet-R introduces recurrent dynamics within each area into the otherwise purely feed-forward network, and CORnet-S is the most complicated (containing the most convolutional layers and including skip connections), aiming to match neural and behavioral data. The three CORnet models have been pre-trained to identify a thousand classes of objects. Parameters of the first three pretrained networks were downloaded from: http://www.vlfeat.org/matconvnet/pretrained/. CORnets were downloaded from: https://github.com/dicarlolab/CORnet. PCA was performed on activation of units in the penultimate layers (IT area in the case of CORnet), and top 50/110PCs were extracted to compare with other models.

##### 5) β-VAE model

We used the standard architecture and optimisation parameters introduced in (Higgins et al., 2017) for training the β-VAE. The encoder consisted of four convolutional layers (32×4×4 stride 2, 32×4×4 stride 2, 64×4×4 stride 2), followed by a 256-d fully connected layer and a 50-d latent representation. The decoder architecture was the reverse of the encoder. We used ReLU activations throughout. The decoder parametrized a Bernoulli distribution. We used Adam optimizer with 1e-4 learning rate and trained the models for 1 mln iterations using batch size of 16, which was enough to achieve convergence. The models were trained to optimize the following disentangling objective:

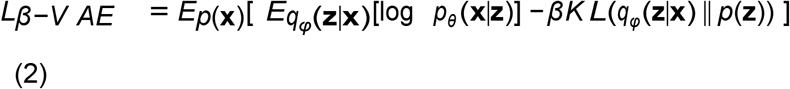

where p(x) is the probability of the image data, q(z|x) is the learnt posterior over the latent units given the data, and p(z) is the unit Gaussian prior with a diagonal covariance matrix.

For the β-VAE model the main hyperparameter of interest that affects the quality of the learnt latent units is the value of β. The β hyperparameter controls the degree of disentangling achieved during training, as well as the intrinsic dimensionality of the learnt latent representation (Higgins et al., 2017). Typically a β>1 is necessary to achieve good disentangling, however the exact value differs for different datasets. Hence, we trained 400 models with different values of β by uniformly sampling 40 values of β in the [0.5, 20] range. Another factor that affects the quality of disentangled representation is the random initialization seed for training the models. Hence, for each β value, we trained 10 models from different random initialization seeds, resulting in the total pool of 400 trained β-VAE.

The recently proposed Unsupervised Disentanglement Ranking (UDR) score (Duan et al., 2020) was used to select 51 model instances with the most disentangled representations (within the top 15% of UDR scores). The UDR score measures the quality of disentanglement achieved by trained β-VAE models (Duan et al., 2020). A set of models were trained using the same hyperparameter setting but with different seeds, and pairwise similarity between the model representations were computed (higher similarity=better disentanglement).

##### 6) Hebbian learning model

This is a biologically plausible model recently proposed to explain view invariance in face cells (Leibo et al., 2017). V1-like features (C1-layer of HMAX model) were extracted from the facial images. PCA was performed on V1-like encodings of a single identity at different head orientations: the *i^th^* PC of the *k^th^* identity is denoted as 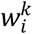. The activation of the *k^th^* unit to a given face is 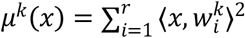, where *x* is the V1-like encoding of that face, and *r* is the number of PCs being used. In our experiment, we used rotated versions of the fitted 3D Morphable Models (from −90° to 90° in 5°increments) as inputs to this model (Figure 2C, 3C), resulting in 2100 such units. PCA was performed on activation of the 2100 units, and the top 50/110PCs were extracted. Since the 3D Morphable Model only fits part of the face and may not provide a satisfactory explanation of neural responses to full faces, we only implemented the Hebbian model on 3D-fits of the original images and compared it to other models under the same condition (Figure 2C, 2D, 3C, 3D).

#### Quantification of explained variance, encoding and decoding errors

To quantify explained variance, a 50-fold cross-validation paradigm was performed: 2100 faces were split into 10 groups of 42 faces. Responses of each neuron to 49 groups of faces were fit by a linear regression model using PCs of a set of features, and the responses of this neuron to the remaining group of 42 faces were predicted using the same linear transform. This process was repeated for all 50 groups, so every single face had a predicted response. Coefficient of determination (R^2^) was used to quantify the percentage of variance explained by the features. Due to noise in the neural data, even the perfect model could not achieve 100% explained variance, therefore the value was further divided by the noise ceiling, which was estimated as 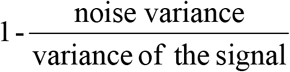. The noise variance was quantified by the mean squared error of neural responses for multiple repetitions of the same stimuli, averaged across all stimuli. Eleven neurons with noisy responses (noise ceiling<10%) were excluded from further analyses, resulting in a population of 148 AM neurons.

For encoding analysis, responses of each neuron were first normalized to zero mean and unit variance. The same procedure was followed to obtain a predicted response for every single face. To quantify prediction accuracy, we examined the predicted responses to individual faces in the space of population responses, and compared this to either the actual response to the face (target) or that to a distractor face. If the angle between the predicted response and distractor response was smaller than that between the predicted response and target response, this was considered as a mistake. Encoding error was quantified as the frequency of mistakes across all pairs of target and distractor faces. Wilcoxon signed rank test was used to determine statistical significance of difference between two models.

For decoding analysis, features of each model dimension were first normalized to zero mean and unit variance. The same procedure used in the encoding analysis was employed, except that the respective roles played by neural responses and model features were reversed.

To quantify the efficacy of the non-overlapping components of two models in predicting neural responses, model features were concatenated together. Explained variance was computed for the combined model, and unique explained variance of a single model was computed by subtracting the explained variance of the other model. Since the combined model has more features, direct comparison will be compromised by an unequal number of parameters. One way to deal with this issue is to use cross validation as in previous analyses--as a higher number of features will be compensated with higher likelihood of overfitting, however, we found the cross-validated explained variances for the combined models were often lower than that for single models, suggesting an overcompensation by overfitting. Therefore we took a different approach: the features of one model were randomly shuffled 50 times and concatenated with the original features of the other model to serve as the baseline, which was compared directly to the combined model. In each shuffle, 50-d features of 2100 faces were randomly permuted across all faces, and the results were averaged across all shuffles.

#### Similarity matrix

Based on the normalized population response, a similarity matrix of correlation coefficients was computed between the population response vectors to each of the n faces. For neural network models, top 50 PCs of activation of units in the penultimate layers of the networks were used to represent faces.

## Acknowledgments

This work was supported by NIH (EY030650-01), the Howard Hughes Medical Institute, and the Chen Center for Systems Neuroscience at Caltech. We are grateful to Nicole Schweers for help with animal training and MingPo Yang for help with implementing the CORnets.

## Author Contributions

L.C. and D.Y.T conceived the project and wrote the paper with the help of all other authors, L.C. performed the experiments and analyzed the data, D.Y.T. supervised the project, B.E and T.V. constructed the 3D Morphable Model used to compare with the neural data.

## Supplementary Information

**Figure S1.**
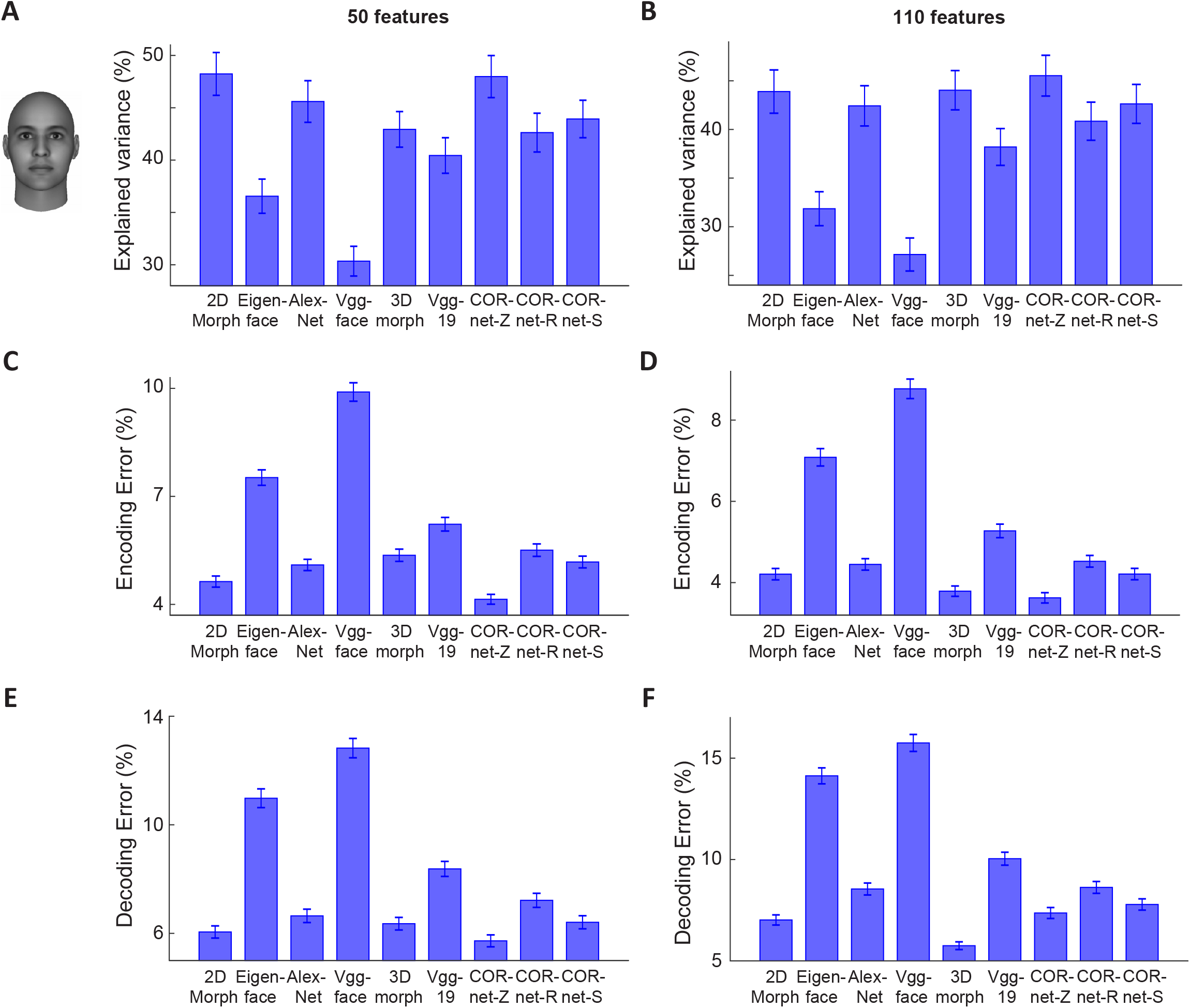
Encoding and decoding analyses using an alternative method of hair removal. Related to Figures 2 and 3. After fitting the original images with the 3D Morphable Model, a mask was created using the fit, and the original image was cropped using the mask. A. Explained variances for 50 features of 9 different models using the cropped image as input. B, save as A, but for 110 features. C and D, same as A and B, but for the encoding errors. E and F, same as A and B, but for the decoding errors.

**Figure S2.**
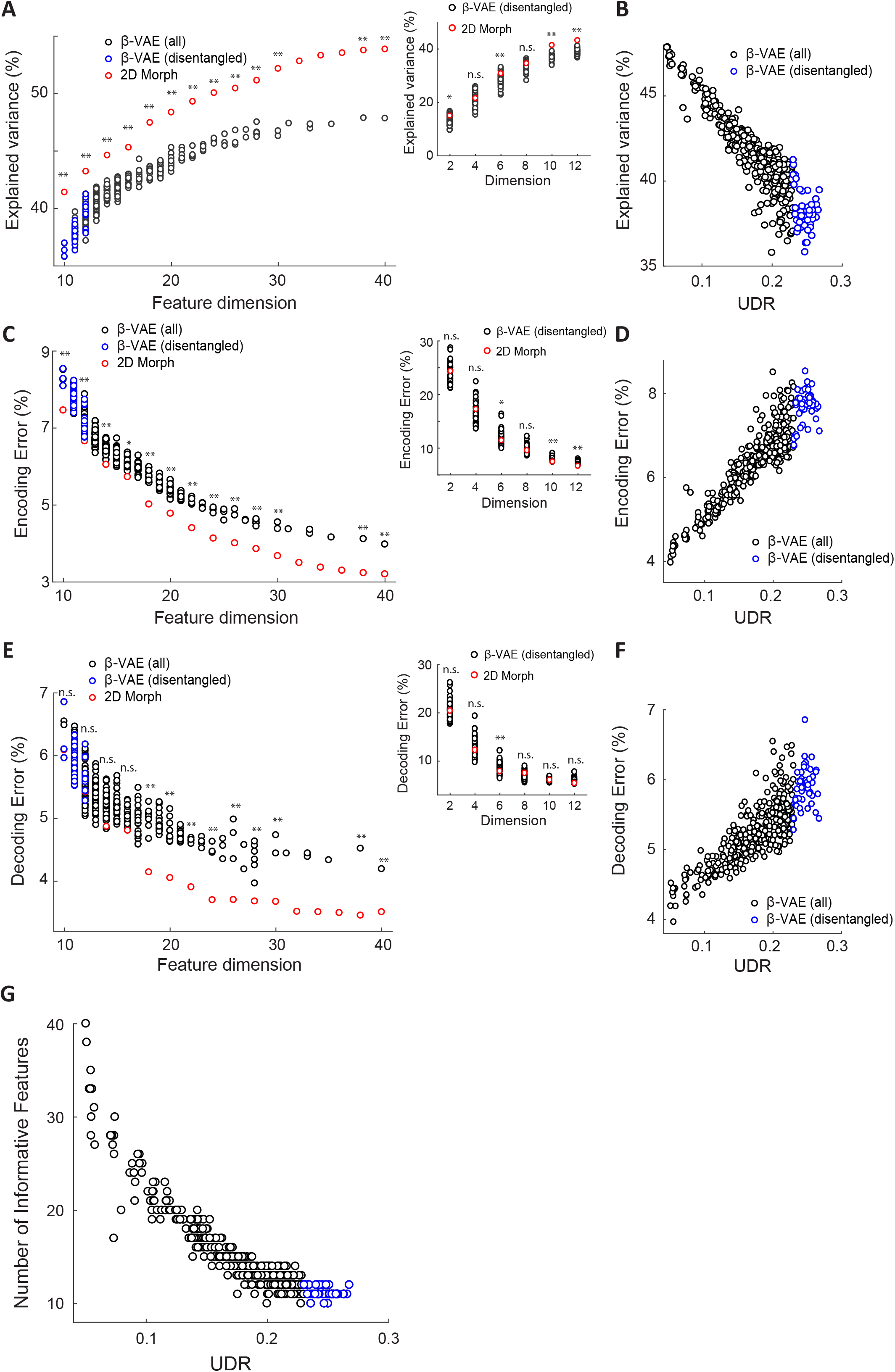
Comparing β-VAEs with 2D Morphable Model at equivalent dimensions. Related to Figures 2 and 3. A, Explained variances for 400 β-VAEs after removing dimensions with variance<0.01 were compared with the 2D Morphable Model at equivalent dimensions (equal number of shape and appearance dimensions were chosen for the 2D Morphable model). Wilcoxon signed-rank test was employed to compare the two models after performing 50-fold cross validation (*=p<0.05; **=p<0.01; n.s.=not significant). Inset, for the 51 most disentangled VAEs, subsets of features explaining the most variance of each model were compared to the 2D Morphable Model at equivalent dimensions (since equal numbers of shape and appearance dimensions were selected for the 2D Morphable model, only even numbers of total dimensions are shown here). B. Explained variances for all 400 β-VAEs are plotted against UDR score. C and D, same as A and B, but for encoding errors. E and F, same as A and B, but for decoding errors. G, Relationship between the number of informative features (variance≥0.01) and UDR score for all 400 β-VAEs.

**Figure S3.**
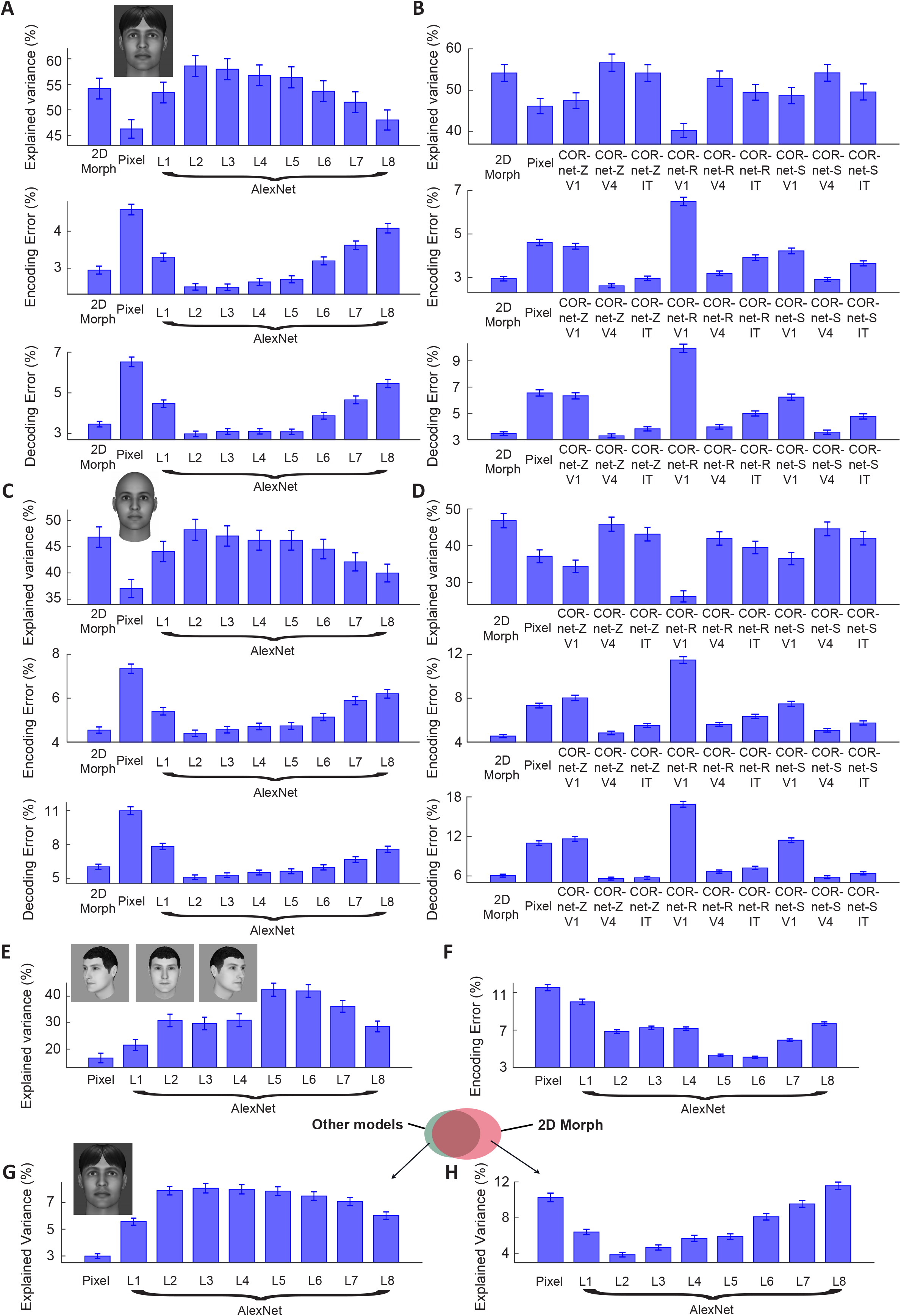
Encoding and decoding performance across different layers of AlexNet and CORnets. Related to Figures 2–4. Inspired by previous works (Yamins et al., 2014; Cadieu et al., 2014), units in the penultimate layer were used to represent different neural network models, we finally examined how each individual layer of AlexNet and CORnets was related to neural responses. Quite surprisingly, we found intermediate layers of those models performed the best, and in particular, 2^nd^ layer (L2) of AlexNet and V4 layer of CORnet-Z even outperformed the 2D Morphable Model (Panels A-D). We next asked whether the same conclusion could be generalized to more complex stimulus sets: previously it was shown that only in face patch AM, neurons achieve full invariance to head orientations--a property unlikely to exist in early layers of CNNs due to the simpler feature selectivity of these layers. We examined the relationship between different layers of AlexNet and AM responses using a stimulus set containing facial images at 8 head orientations (Panels E and F). We found in this case, AlexNet L2 lost its advantage, with L5/6 performing the best. Given that AlexNet L2 cannot explain neural responses to the stimulus set with different head orientations, it is clearly not a viable candidate for explaining face patch activity. Finally, we asked how much unique variance each layer of AlexNet explained compared to 2D Morph by repeating the analysis in Figure 4A on individual layers of AlexNet, and found that the amount of unique variance of AlexNet L2 is similar to that of other intermediate layers (L3 to L6, Panel G). We also asked the converse, how much unique variance is explained by 2D morph compared to each of the layers of AlexNet (Panel H). Here, we found what makes L2 distinct: the 2D Morph Model explains the least unique variance when compared to L2 activation. A, Explained variances, encoding errors and decoding errors of the 2D Morphable Model and multiple layers of AlexNet for images after background removal (cf. Figure 2B). Pixel=input to the network. B, Same as A, but for CORnets. C and D, same as A and B, but for images fit by the 3D Morphable Model. E and F, Explained variances and encoding errors of a stimulus set containing 28 identities with 6-8 head orientations for multiple layers of AlexNet. Due to the small size of the stimulus set (n=220), 15 features were used instead of 50 to avoid overfitting, but the trend remained the same with 50 features. [Note that the images shown here are synthetically generated faces that serves as stand-ins for the actual example images in order to satisfy bioRxiv’s policy on the use of images of human faces.] G and H, same as Figure 4A and B, but for multiple layers of AlexNet.

**Figure S4.**
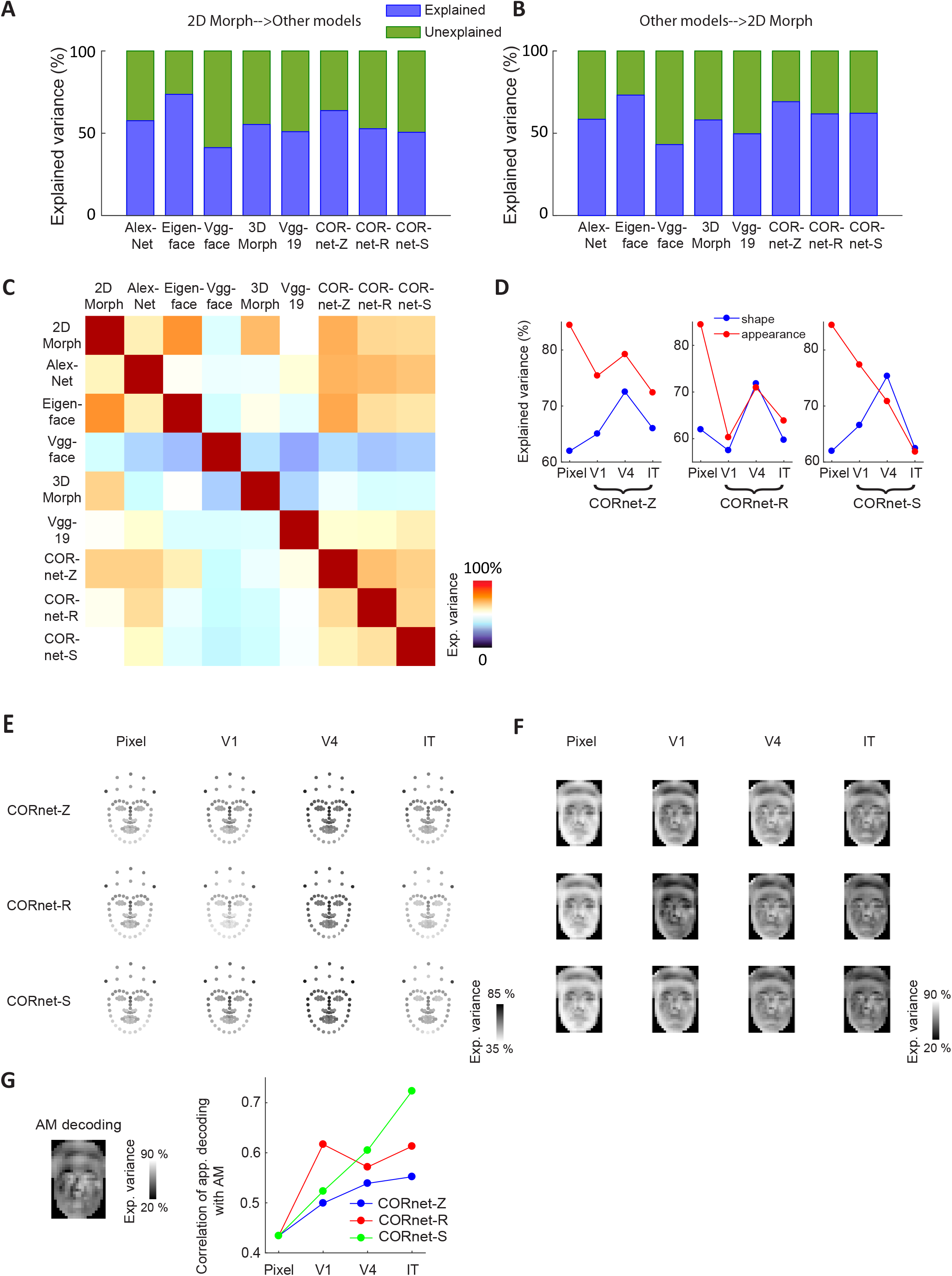
Direct comparison between feature spaces spanned by different models. Related to Figure 4. A, Quantification of how much variance of features in one model could be explained by features of the 2D Morphable Model (similar to Figure 4B, but using feature values directly instead of neural responses). B, Same as A, but with features of the 2D Morphable Model being fitted by the other models. C, For each pair of two different models (X,Y), features of model X were fit by features of model Y, both using 50 feature dimensions (PCs). Explained variances were then averaged across the 50 PCs of model X, weighted by the variance of the original features explained by each PC. The averaged explained variances of all model pairs were then color-coded and plotted as a matrix, with its rows representing model Xs, and columns representing model Ys. D, Shape and shape-free appearance features for the 2D Morphabale Model were separately fit by 4 different layers (“Pixel”=input to the network) of three CORnet models. The V4 layer performed best for shape features, while the input layer performed best for shape-free appearance features. E, Decoding performance of all landmarks for different layers of CORnets. F, Decoding performance of shape-appearance descriptors for different layers of CORnets. All images were first morphed to the average shape. Pixel intensities of the shape-free image were averaged within multiple grids across the image (29*20 grids in total), and then fit with model features. G, Correlation of the decoding maps between CORnets and neural data.

**Figure S5.**
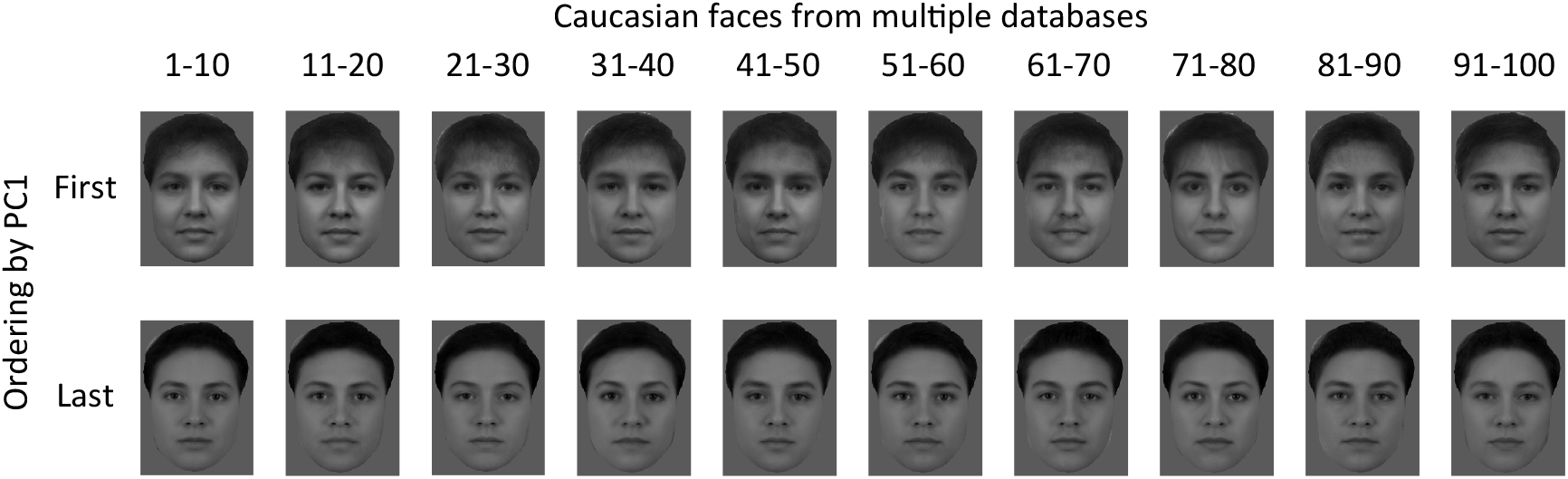
Comparison between VGG-face and AlexNet for Caucasian faces. Related to Figure 5. Same as Figure 5B, but for 748 Caucasian faces. To attenuate the influence of diverse image backgrounds in multiple databases, we removed the background before presenting images to the networks (cf. Figure 2B). [Note that images shown here are not actual faces of any individuals, but the average images of 10 faces after being morphed to the average shape, using the same algorithm as the 2D Morphable Model.]

